# Effect of SFX-01 on proliferation of human glioblastoma cell lines

**DOI:** 10.1101/2021.09.14.459936

**Authors:** Euphemia Leung, Edwina Wright, Bruce C. Baguley

**Affiliations:** Auckland Cancer Society Research Centre, University of Auckland, Auckland, New Zealand; Maurice Wilkins Centre for Molecular Biodiscovery, University of Auckland, Auckland, New Zealand; Evgen Pharma PLC, Alderley Park, Cheshire, United Kingdom

**Keywords:** Sulforaphane, SFN, SFX-01, α-cyclodextrin, glioblastoma, spheroids, cell culture

## Abstract

Glioblastoma (GBM) is the most aggressive and lethal of brain tumours and there is a clear need for novel therapies. SFX-01, a stabilised formu of sulforaphane (SFN) that is complexed with α-cyclodextrin, is being investigated as a potential anticancer drug. And as part of this investigation we have determined its effect on the proliferation of a series of tumour cell lines. Growth-inhibitory concentrations (IC_50_ values) of SFX-01 and reference SFN were highly correlated but showed a divergence at higher drug concentrations that is likely to reflect drug binding to α-cyclodextrin complexes. Because SFN is redox-sensitive, we also compared the effects of these drugs on four GBM lines that had been derived and cultured under physiological oxygen (5%) conditions. Comparisons were also made using a three-dimensional spheroid model, used to mimic the *in vivo* state of glioblastomas and to simulate barriers to drug delivery *in vivo*. SFX-01 and reference SFN and inhibited proliferation of four GBM cell lines in monolayers as well as in spheroids; α-cyclodextrin alone had no significant effect. Our results are consistent with the hypothesis that SFN is liberated from SFX-01 at concentrations sufficient to inhibit cell growth. We envision that SFX-01 will have improved pharmacokinetic properties *in vivo*, warranting further pre-clinical investigation and clinical development.

## Introduction

While the outlook for many types of cancer types has improved with therapeutic advancements over past years, that for glioblastoma (GBM) remains poor with only 5% of patients surviving 5 years post-diagnosis (https://www.curebraincancer.org.au/page/8/facts-stats). Current therapy generally comprises surgery, followed by radiotherapy and chemotherapy with temozolomide, with a median survival of less than 16 months.^1^ There is a need for novel approaches to therapeutic agents.

SFN is an isothiocyanate compound derived from cruciferous vegetables, particularly broccoli and broccoli sprouts, and is an agent with potent anti-oxidant and anti-inflammatory activity.^2^ SFN suppresses the growth of glioblastoma cells, glioblastoma stem cell–like spheroids, that targets several apoptosis and cell survival pathways.^3^ However, clinical application has been hindered by its physicochemical and biological instability.^4^ To improve the stability of SFN, SFX-01 (a fully synthetic SFN complexed with α-cyclodextrin) has been developed for clinical use under the European patent EP2854861 (<u>Cyclodextrins | European Medicines Agency (europa.eu)</u> - https://www.ema.europa.eu/en/cyclodextrins).

*In vitro* assessment of new drugs requires consideration of the physicochemical properties, and SFN is oxidation-sensitive. While we have compared the growth inhibitory concentrations of SFX-01 and reference SFN under standard conditions, we have also compared them under physiological oxygen (5%) conditions, chosen to minimise the effects of oxygen, for a series of four cell lines that were also derived under these conditions. Assessment should also recognise the contribution of cancer stem cells (CSCs), which are involved in both tumour initiation and growth, as well as modulating resistance to chemotherapy and radiotherapy ^5^ limiting long-term survival. Spheroid formation has been proposed as a robust, independent predictor of glioma tumour progression^6^ and the presence of glioma stem cells (GSCs), which are enriched in spheroids, may be connected to the drug resistance of GBM due to their enhanced antioxidant defence^7^ and elevated DNA repair capacity.^8^

In this report we firstly compared the growth inhibitory capacity of SFN from SFX-01 versus reference SFN using a series of tumour cell lines. Secondly we compared their effects against four low-passage GBM cell lines that had been generated and grown under physiological oxygen conditions. Thirdly we compared the drug effects on these four cell lines grown either in monolayer cultures or in 3-dimensional spheroid cultures. Fourthly we determined whether α-cyclodextrin itself affected the proliferation of the four GBM cell lines.

## Materials and Methods

### Chemicals

Sulforaphane reference material (catalogue number S4441) and α-cyclodextrin (Cavamax-W6) (catalogue number 778788) were purchased from Sigma-Aldrich (New Zealand). SFX-01 was a gift from Evgen Pharma PLC (https://evgen.com/).

### Cell lines

Four NZ brain GBM cell lines NZB9, NZB11, NZB13 and NZB18 had been generated at the Auckland Cancer Society Research Centre ^9^ under 5% oxygen to mimic, as far as possible, physiological conditions. As described previously, ^10^ cell lines originated from tumour tissue taken from four patients undergoing surgery for brain cancer. Tumour tissue was sent to the histopathologist immediately after surgery. Formal consent had been obtained from all patients, using guidelines approved by the Northern A Health and Disability Ethics Committee.

Solid tumour specimens were disaggregated either immediately or after overnight storage at 4°C. Normal, adipose, or grossly necrotic material was removed and the tumour tissue was minced finely using crossed scalpels. Tissue was reduced to small clumps by passage through a 0.65-mm stainless steel sieve. The size of the aggregates varied greatly (aggregates of 5–100 cells). Material containing larger aggregates was pipetted into tubes, collected by low-speed centrifugation to remove blood cells, necrotic material, and debris (30 × g, 2min), and then washed twice (30 × g, 2min) in growth medium. Preparations were monitored by phase contrast microscopy, and cytospins of cell suspensions were stained by haematoxylin/eosin and examined by a pathologist to ensure that they contained tumour cells. Cultures were set up in growth medium supplemented with 5% FBS under an atmosphere of 5% O_2_, 5% CO_2_, and 90% N_2_ in a Tri-Gas Forma incubator. Tissue culture plates or flasks had previously been coated with a thin layer of agarose to prevent the growth of fibroblasts. ^11^ Cells were cultured in either αMEM (12,000,063, Thermo-Fisher Scientific) supplemented with 5% v/v foetal bovine serum (FBS; Moregate Biotech, Hamilton, New Zealand) for monolayers or cultured in DMEM:F12 (1:1, ThermoFisher Scientific) supplemented with B-27 (17504044,, Thermo-Fisher Scientific) for spheroids. Early passage cell lines (< passage 25) used in this study were developed in this laboratory. All cell lines were tested negative for mycoplasma contamination.

Human brain cancer U87mg and U251 cell lines, melanoma A375 cell lines, H460 lung cell lines, triple-negative breast cancer (TNBC) MDA-MB-231 cell line and HCT116 colon cancer cell lines were purchased from the American Type Culture Collection (ATCC). These cell lines were passaged in αMEM supplemented with 5% foetal bovine serum (FBS), and insulin/transferrin/selenium supplement (Roche) under ambient (21%) oxygen conditions without antibiotics for less than 3 months from frozen stocks confirmed to be Mycoplasma-free.

### 3D spheroid cell culture

On day 0, 1 × 10^4^ cells/well were seeded into ultra-low attachment round bottom 96-well plates (7007, Corning, Kennebunk, ME) in 50 μL DMEM:F12 supplemented with B-27. The plates were centrifuged at 1000×g for 10 min and placed in the cell culture incubator to allow establishment of spheroids. On day 1, 50 μL of fresh DMEM:F12 supplemented with B-27 was added to the plates, and 20 μL of DMEM:F12 + B-27 containing SFX-01 or SFN at the desired final concentration was added to the spheroid plates with the maximum concentration of 50 μM. On day 5, ^3^H-thymidine was added to the cultures and thymidine incorporation was detected following a 16 h overnight incubation.

### ^3^H-thymidine incorporation assay

Proliferation was measured using a thymidine incorporation assay, as described previously.^12^ Briefly, cells were seeded (1–3 × 10^3^ per well) in 96-well plates in the presence of varying concentrations of inhibitor for 3 or 5 days, as indicated. ^3^H-thymidine (0.04 μCi per well for two dimensional (2D) monolayer culture or 0.08 μCi per well for three dimensional (3D) spheroid culture) was added (6 h for 2D monolayer culture or 16 h for 3D spheroid culture) prior to harvest, cells were then harvested on glass fibre filters using an automated Tom-Tec harvester. Filters were incubated with Betaplate Scint and thymidine incorporation measured in a Trilux/Betaplate counter. Effects of inhibitors on ^3^H-thymidine incorporation into DNA were determined relative to the control cells incubated with water or DMSO (vehicle treated) samples.

### Data analysis

*T*-test was used for comparison between two groups. Correlation analysis was performed with Spearman’s rank order correlation coefficient (R) and statistical significance (P) using Prism (version 8.0.2, GraphPad Software, Inc.). Values of *p* < 0.05 were considered to be statistically significant.

## Results

### Comparison of the ability of SFN and SFX-01 to reduce proliferation in cancer cell lines grown in monolayer culture

The effect of SFN and SFX-01 on cell proliferation was examined in a panel of cancer cell lines grown as monolayers (Figure 1). A significant correlation was observed between sensitivity of the cell lines to SFX-01 and SFN (Table 1). However, the two sets of growth inhibitory concentrations (IC_50_ values) diverged at higher values, with higher values SFX-01. The correlation between the IC_50_ ratios (SFX-01/SFN) and the IC_50_ values for SFX-01 was found to be highly significant (*r* = 0.96; *p* < 0.001), suggesting that a proportion of SFX-01 was sequestered and not available to the cells.

**Figure 1.**
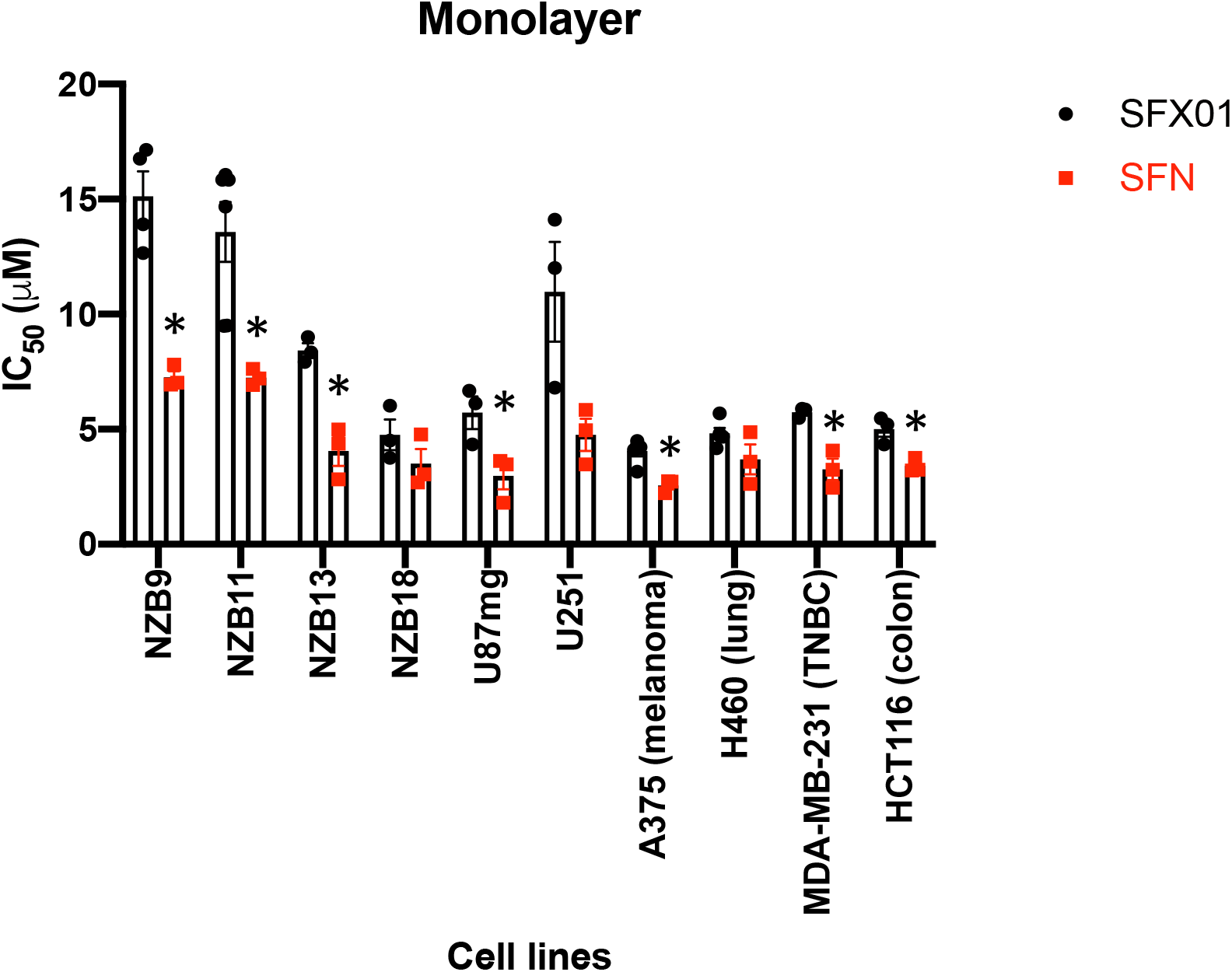
Comparison of the effect of SFX-01 and SFN in a panel of cancer cell lines. Brain cancer cell lines NZB9, NZB11, NZB13, NZB18, U87mg, U251, melanoma cell line A375, lung cancer cell line H460, triple-negative breast cancer cell line MDA-MB-231 and colon cancer cell line HCT116 were exposed to the compounds for 3 days and inhibitory effect was measured by thymidine uptake. *, *p* > 0.05, multiple unpaired T test.

**Table 1.** The IC_50_ values between SFN and SFX-01 were comparable (Spearman Rank Order Correlation with *r* = 0.781, *p* (2-tailed) = 0.0076) in the cell lines grown as monolayer culture.

|  | SFX-01 | SFN |
| --- | --- | --- |
|  | IC<sub>50</sub> (μM) + SE |  |
| NZB9 | 15.1 ± 1.1 | 7.3 ± 0.3 |
| NZB11 | 13.6 ± 1.3 | 7.2 ± 0.2 |
| NZB13 | 8.4 ± 0.3 | 4.1 ± 0.6 |
| NZB18 | 4.8 ± 0.7 | 3.5 ± 0.6 |
| U87mg | 5.7 ± 0.7 | 3.0 ± 0.6 |
| U251 | 11.0 ± 2.2 | 4.8 ± 0.7 |
| A375 (melanoma) | 4.1 ± 0.2 | 2.6 ± 0.2 |
| H460 (lung) | 4.8 ± 0.2 | 3.7 ± 0.7 |
| MDA-MB-231 (TNBC) | 5.7 ± 0.1 | 3.3 ± 0.5 |
| HCT116 (colon) | 5.0 ± 0.3 | 3.4 ± 0.2 |

### Effects of SFN and SFX-01 on proliferation of GBM cell lines grown as neurospheres

The anti-proliferative effects of SFN and SFX-01 in NZB9, NZB11, NZB13 and NZB18 cell lines are compared in Table 2 and Figure 2. Although spheroid formation was apparent in NZB18, cell proliferation ceased by Day 5 and therefore proliferation could not be measured by thymidine uptake. There was no significant difference in IC_50_ values between SFN and SFX-01 in the three NZ GBM cell lines tested (Table 2).

**Table 2.** IC_50_ values for SFN and SFX-01 in spheroid culture There was no significant difference (*p* > 0.05, paired T-test) in the three NZ GBM cell lines.

|  | SFX-01 | SFN |
| --- | --- | --- |
|  | IC<sub>50</sub> (μM) + SE |  |
| NZB9 | 37.0 ± 2.0 | 40.5 ± 3.7 |
| NZB11 | 36.3 ± 7.1 | 30.9 ± 5.6 |
| NZB13 | 25.5 ± 3.7 | 16.9 ± 2.6 |

**Figure 2.**
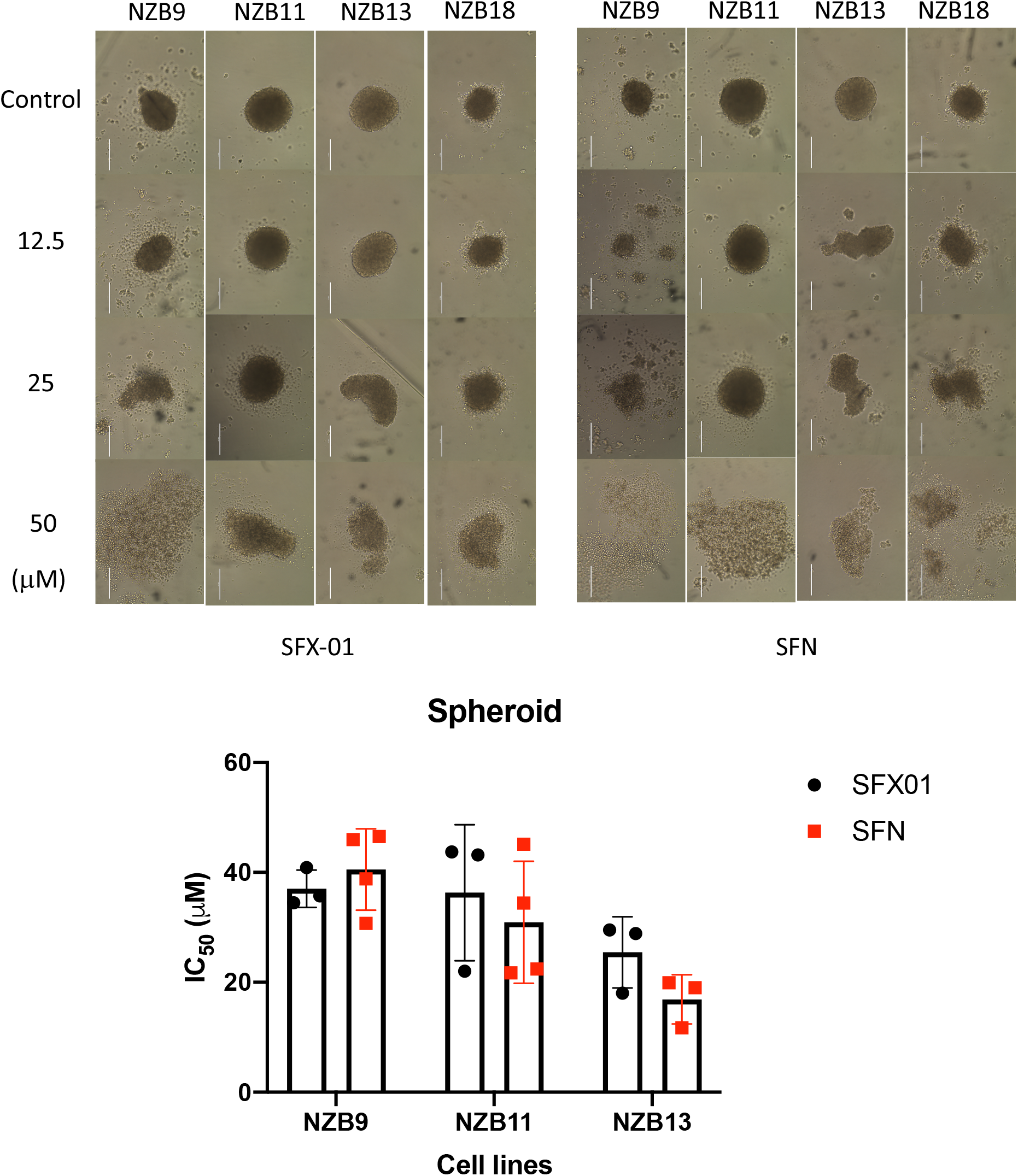
(Top) Images of four NZ brain cancer spheroid cell cultures captured on day 5 (scale bar = 0.4 mm). (Bottom) IC_50_ values measured by thymidine incorporation following 5 days of treatment with SFX-01 and SFN, respectively (mean ± SEM, n = 3-4).

### Effect of α-cyclodextrin on proliferation of four New Zealand brain cancer cell lines

α-cyclodextrin is used to stabilise SFN in the SFX-01 complex. The four NZ brain cancer cell lines were grown as monolayer and no significant reduction in proliferation was observed when incubated with α-cyclodextrin (Figure 3).

**Figure 3.**
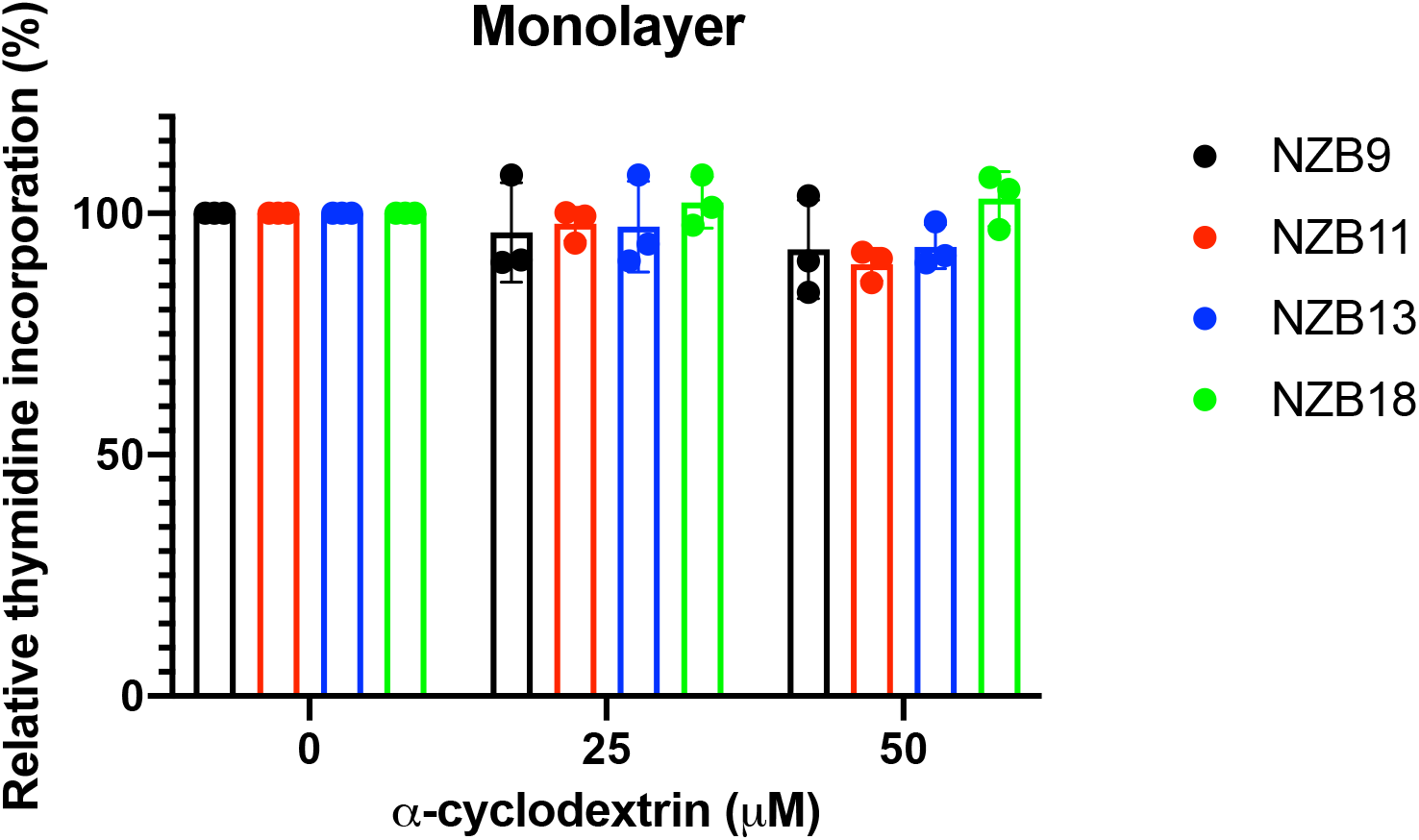
Effect of α-cyclodextrin on the proliferation of the four New Zealand brain cancer cell lines.

## Conclusions

The results show that IC_50_ values for SFN and SFX-01 for each of the cell lines are correlated but the values for SFX-01 are higher for the NZB18, U251 and H460 cell lines, which show higher IC_50_ values. The most likely explanation for this behaviour is that SFX-01 binding to α-cyclodextrin slightly decreases its bioavailability *in vitro*, as implicated by studies of cyclodextrin binding to other drugs.^13^ Thus, binding of SFN to α-cyclodextrin reduces its bioavailability in these culture systems, particularly at SFX-01 concentrations higher than 5 μM. However, only SFN released by SFX-01 is expected to be present in the systemic circulation and cross the blood brain barrier. It is because the carrier (α-cyclodextrin) is thought to remain in the gastrointestinal tract.^14^ This may simply be a feature of *in vitro* evaluations of the agent.

The IC_50_ values of the four low-passage GBM cell are generally higher than those of the other lines but there is a considerable overlap. Thus, it cannot be concluded that the physiological oxygen concentrations leads to altered IC_50_ values lines that had been generated and grown under physiological oxygen conditions. However, it is clear that when these cell lines are grown as monolayer cultures they are more sensitive to SFN and SFX-01 than they are 3-dimensional spheroid cultures. The reduced sensitivity of spheroids is likely to result from cytokinetic effects (a decreased proportion of cells undergoing DNA replication) as well as from drug binding to α-cyclodextrin. The results also demonstrate that the presence of α-cyclodextrin itself is unlikely to change the response of the four GBM cell lines.

## Acknowledgements

The authors thank Drs. Huw Jones, Nick Mallard and Bruno Simoes (orcid: 0000-0003-1253-6657) for their helpful discussions.

